# Natural and synthetic polyamines as organic counterions for multi-crossover DNA nanostructures

**DOI:** 10.64898/2026.11.24.768944

**Authors:** Bharath Raj Madhanagopal, Gwenyth Gallagher, Nada Kabbara, Vinod Morya, Mengyao Zheng, Ken Halvorsen, Arun Richard Chandrasekaran

## Abstract

Assembly of DNA nanostructures requires positive counterions to shield the negative charge repulsion between the strands in the assembly. While several counterions have been explored for DNA nanostructure assembly, polyamines are particularly attractive candidates because they could support intracellular assembly of DNA nanostructures. However, a systematic analysis of natural and synthetic polyamines for DNA nanostructure assembly is currently lacking. In this study, we demonstrate the assembly of three model DNA nanostructures of the same size but with different number and topology of crossovers using three naturally occurring and two synthetic polyamines. The model DNA nanostructures are a double crossover motif with two antiparallel crossovers (DX-A), a double crossover motif with two parallel crossovers (DX-P), and a paranemic crossover with six parallel crossovers (PX). We successfully assembled all nanostructures, and relative assembly yields depended on the concentration and the type of polyamine and the type of DNA nanostructure. The structures exhibited comparable conformations but displayed slightly lower thermal stabilities compared to their Mg^2+^ assembled counterparts. We also demonstrate hierarchical assembly of DX tiles into periodic 2D lattices using the polyamines. Our results shed light on how polyamines differentially impact DNA nanostructures with distinct topological features and different crossover densities. Polyamines can serve as organic biocompatible counterions for the assembly of simple DNA motifs as well as larger arrays with implications for intracellular assembly and biological applications of DNA nanostructures.

---

The programmable assembly of DNA strands has been used to produce a broad range of designer DNA nanostructures. These nanoscale structures have found several applications across fields such as diagnostics, drug delivery, materials science, and computation. Although Mg^2+^ ions are typically used in the assembly of DNA nanostructures, they are not compatible for all applications and exploration of Mg^2+^ alternatives is a key focus area in DNA nanotechnology.^1^ Several alternative counterions have been shown to work in a structure-dependent and application-dependent manner.^2–6^ In addition, alternatives to metal ions have also been explored, such as the use of organic solvents,^7^ peptides,^8^ and cationic amino acids.^9^

Polyamines are a class of positively charged biomolecules present in all living organisms that act as organic counterions with essential functions such as chromatin condensation, cell growth, and cell signaling.^10,11^ Polyamines bind to DNA primarily through electrostatic interactions due to their multiple positive charges on their amine groups, neutralize the negatively charged phosphate backbone of DNA, and facilitate DNA self-assembly.^12,13^ While the typical monovalent and divalent metal cations, such as Na^+^ and Mg^2+^ act as concentrated point charges, the positive charges in polyamines are separated by hydrocarbon chains. The spacing between the amine groups allows polyamine molecules to establish multiple points of contact at the backbone and the major and minor grooves of the double helix.^14^ For example, spermine, spermidine, and putrescine, polyamines present in eukaryotic cells, contain 4, 3, and 2 ionizable amine groups respectively, and bind to DNA to varying affinities.^15^ Some polyamines can also act as molecular bridges and connect adjacent DNA double helices. This neutralizes the inter-helix electrostatic repulsion and allows for the DNA to fold and form highly ordered condensed units. With a diversity of charge density, propensity for DNA binding, and biocompatibility, polyamines are suitable substitutes for metal counterions in DNA self-assembly, especially for biological applications.

Diverse nanostructures such as tetrahedra,^16,17^ nanoprism,^18^ and DNA origami^19^ have been assembled using polyamines as the counterion and are shown to be promising DNA nanocarriers. For example, polyamine-stabilized DNA tetrahedra displayed higher cell uptake efficiency compared to Mg^2+^-stabilized nanostructure and delivered siRNA.^17^ Spermidine improves the stability of DNA origami structures in cell lysates and enhances the stability of the nanostructures under electric field pulses, increasing their potential as electrotransfection agents.^19^ The use of spermidine in DNA nanostructures also increases the anti-inflammatory properties of the DNA nanoplatforms such as tetrahedra and DNA hydrogels.^17,20^ In addition, an azide-bearing spermine was used to fold DNA origami in the absence of Mg^2+^ and provide reactive groups for further functionalization.^21^

Multi-crossover DNA tiles are basic structural units used in tile-based assemblies of DNA. Similar to DNA origami and other smaller discrete nanostructures, the assembly of multi-crossover tiles and their hierarchical assemblies depends on effective electrostatic shielding of the negative charges of the DNA backbone. Although polyamines have been used to assemble tetrahedra, prisms, and DNA origami structures, a systematic study into their ability to assemble multi-crossover DNA tiles has been investigated only by computational methods^22,23^ and has not been experimentally demonstrated. Here, we demonstrate the assembly of three types of DNA multi-crossover motifs using both naturally occurring and synthetic polyamines. We further demonstrate successful assembly of a DNA tile-based 2D array using these polyamines.

In this study, we chose spermine, spermidine, putrescine, N-N’-bis(3-aminopropyl)ethylenediamine (BAPD), and tris(2-aminoethyl)amine (TAEA) as model polyamines (**Figure 1**). While spermine, spermidine, and putrescine are naturally occurring polyamines present in most eukaryotic cells, BAPD and TAEA are synthetic polyamines generally used as chelating agents and building blocks in synthetic chemistry.^24,25^ We chose these polyamines as they are inexpensive and easily available. BAPD is similar to spermine in the number of primary and secondary amines but differs in the number of carbons separating the two secondary amine groups. While the natural polyamines and BAPD contain a linear backbone, TAEA represents branched polyamines.

**Figure 1.**
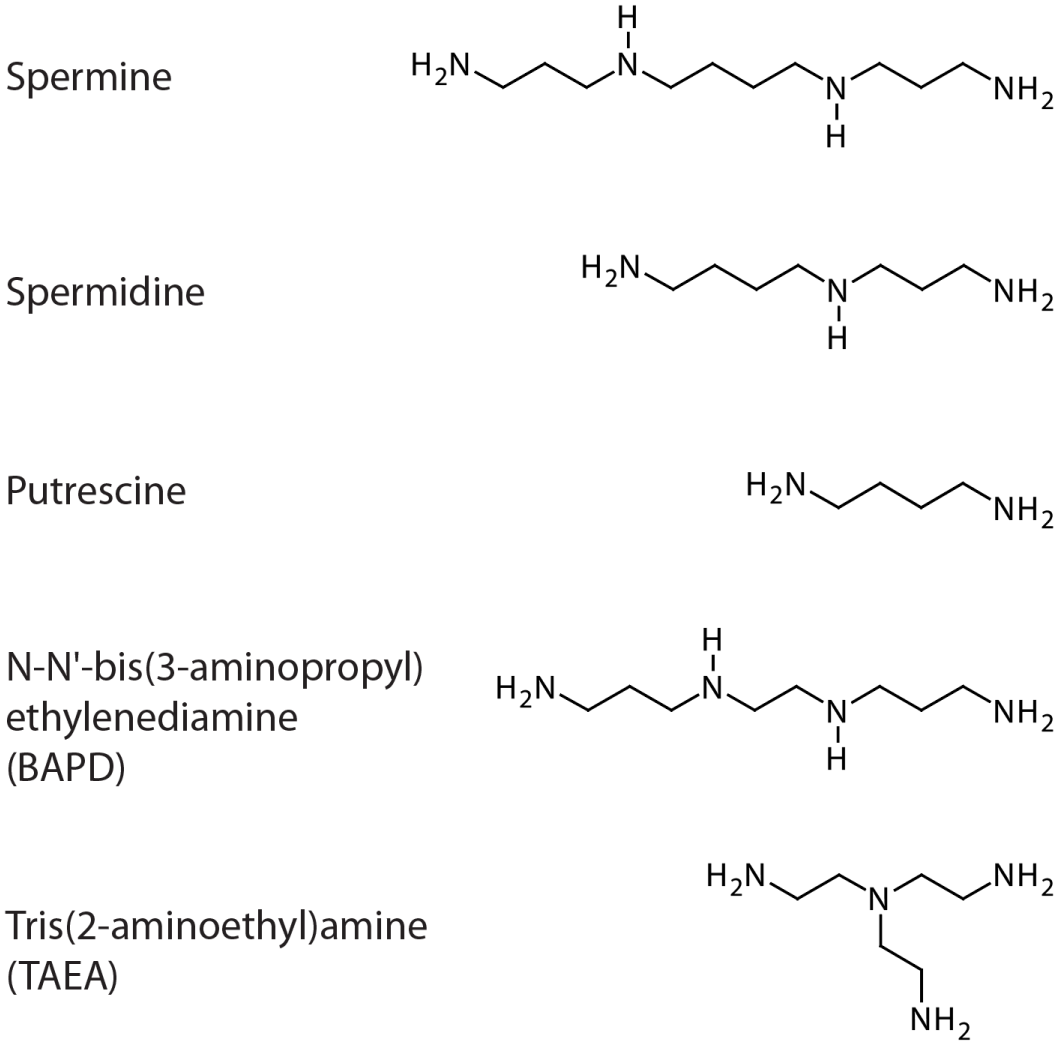
Chemical structure of polyamines used in this study. Spermine, spermidine, and putrescine are naturally occurring polyamines. BAPD and TAEA are synthetic polyamines.

To study the effects of polyamines on the assembly of multi-crossover DNA motifs, we chose the double crossover (DX) DNA motif with antiparallel crossovers (DX-A),^26^ DX DNA motif with parallel crossovers (DX-P),^26^ and a paranemic crossover (PX) DNA motif^27^ as model structures (**Figure 2a, Figure S1 and Table S1**). All three structures contain the same number of putative base pairs (76 bp) and consist of 4 component strands, each with a unique sequence. The GC contents of DX-A, DX-P, and PX are 56.7%, 44.7%, and 44.7%, respectively. All three structures contain two double helical domains. In both DX-A and DX-P, the double helical domains are linked by two crossovers, but they differ in the crossover topology. While the two crossovers in DX-A are antiparallel, they are parallel in DX-P.^26^ On the other hand, the two duplex domains in PX are linked by six parallel crossovers instead of two. The tightly packed PX structure with consecutive crossovers has shown exceptional biostability and limited enzyme interaction compared to duplexes and DX motifs.^28,29^ With these models, we investigated whether the ability of the polyamines to fold DNA and assemble them into nanostructures is influenced by the hybridization topology and crossover density.

**Figure 2.**
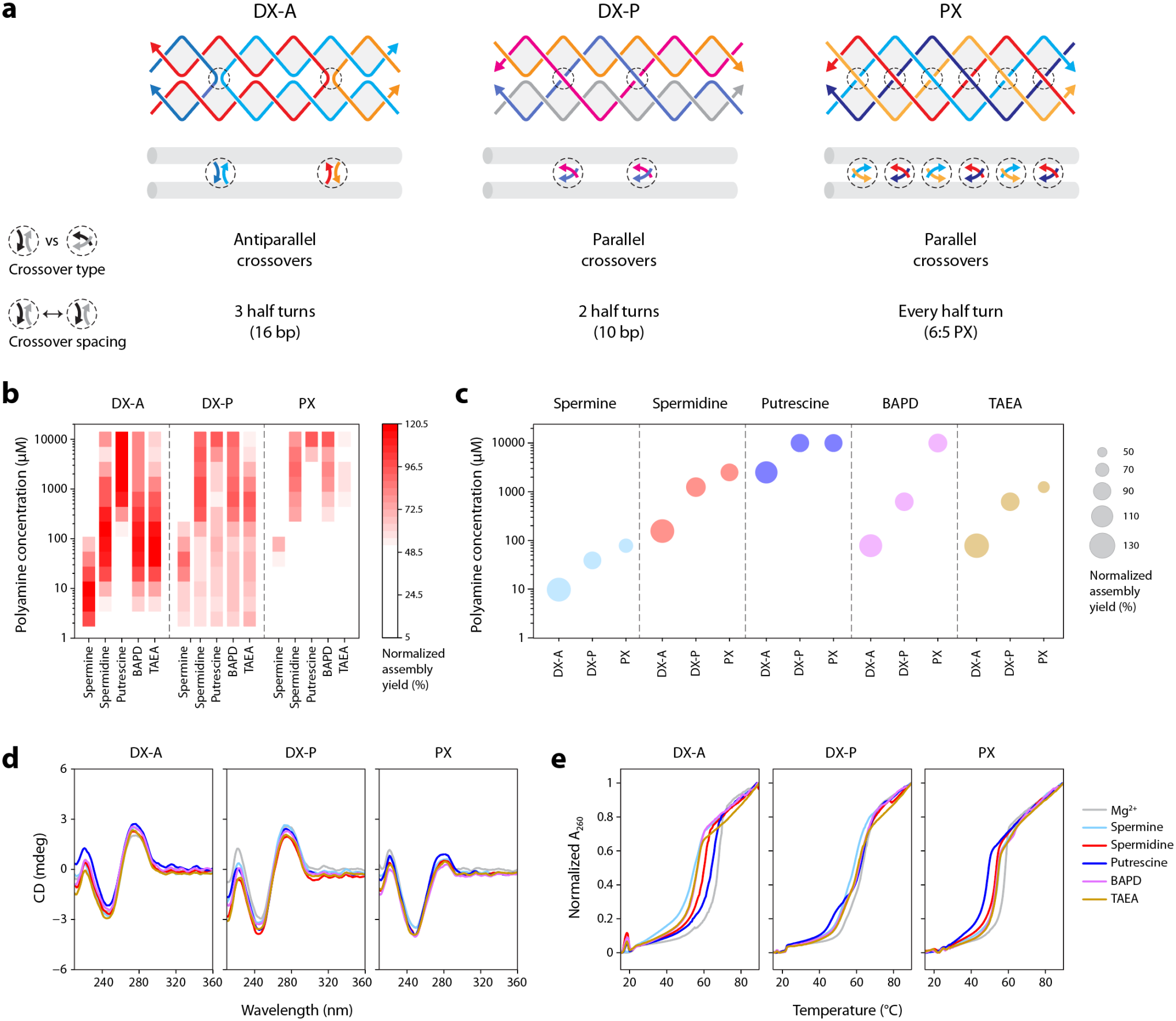
Self-assembly of multi-crossover motifs in polyamines. (a) Schematic of the motifs tested: double crossover motif with antiparallel crossovers (DX-A), parallel crossovers (DX-P) and a paranemic crossover (PX) DNA motif. (b) Assembly yields of the motifs in different polyamines normalized to assembly yields in Mg^2+^ ions. (c) Concentration of polyamines required for the highest normalized assembly yields of the three nanostructures. (d) CD spectra of motifs. (e) UV-thermal melting curves of DX-A, DX-B, and PX.

First, we compared the assembly yields of DX-A, DX-P, and PX in the five different polyamines over a range of polyamine concentrations. We assembled all the structures by mixing the four component strands in stoichiometric amounts in 1× tris acetate-EDTA (1× TAE) buffer (pH 8) supplemented with 12.5 mM magnesium acetate or different concentrations of the polyamines. We annealed the mixture over a thermal gradient from 90 °C to 4 °C of ∼2 hours. Expecting that the concentration range might be different for different DNA nanostructures and polyamines, we chose to screen a large range from 2.4 μM to 10 mM. We ran the assembled nanostructures on non-denaturing polyacrylamide gels and calculated the relative assembly yields of structures in the polyamines compared to the structures assembled using 12.5 mM magnesium acetate.

In general, the polyamines tested in this study appeared to perform well at different concentration ranges. Spermine showed higher assembly at 2-75 μM while the minimum concentration of spermidine, BAPD, and TAEA needed for efficient assembly was between 75 and 625 μM (**Figure 2b** and **Figure S2**). Compared to other polyamines, putrescine was less efficient at assembling the nanostructures at lower concentrations and displayed high assembly yields only at millimolar concentrations. Specifically, DX-A formed efficiently with relative assembly yields comparable to Mg^2+^ at lower concentrations of spermine (∼2-10 μM) compared to DX-P (∼40 μM) and PX (∼75 μM), with the assembly yields declining at higher concentrations. In spermidine, highest relative assembly yields for DX-A was observed at ∼75–150 μM while highest yields for DX-P and PX were seen at 625 μM and above. Putrescine showed efficient assembly of DX-A only above 625 μM and remained high up to 10 mM polyamines (highest concentration tested). However, efficient assembly of DX-P and PX was observed only at 10 mM putrescine. The trend displayed by BAPD was similar to spermidine, with PX requiring slightly higher concentrations of the polyamine. TAEA showed the highest DX-A assembly yield at 75 μM. The efficiency of DX-P assembly peaked at 625 μM and declined at higher concentrations. On the other hand, PX assembly by TAEA was inefficient with only a highest of ∼60% yield at concentrations above 625 μM. The decline in the assembly of the nanostructures seen at higher concentrations of spermine and spermidine is consistent with previous reports and is probably due to aggregation.^16,30^

The polyamines tested here differ in the number of amine groups, length of the aliphatic chains separating them, and degree of backbone branching. Since the interaction of polyamines with DNA is influenced by these aspects of their chemical structure, these features may also influence the efficiency of the polyamines to assemble DNA nanostructures.^31–33^ A comparison of the concentration of polyamines required for highest assembly yield showed a structure-specific trend (**Figure 2c**). Between the antiparallel and parallel crossovers, DX-A required lower polyamine concentrations for efficient assembly compared to DX-P. Comparing DX-P and PX, the polyamine concentration required for assembly is higher when the crossover density increases, with PX requiring higher polyamine concentrations across all the polyamines. Notably, polyamines also allow isothermal assembly of these motifs at the physiological temperature of 37 °C (**Figure S3**). These results show that polyamines allow successful assembly of DNA motifs at constant moderate temperatures similar to monovalent and divalent ions previously reported,^2,6,34^ with only spermidine used for isothermal assembly^18^ prior to this study.

Next, we performed circular dichroism analysis of the nanostructures prepared using the best assembly conditions for each polyamine to determine whether polyamines induced any conformational changes to the structure (**Figure 2d**). All three structures showed a positive band at ∼275 nm. DX-A and DX-P showed a negative band at ∼245 nm, and this band was slightly shifted to the right to ∼250 nm in PX. The positive band of PX at ∼275–280 nm was less intense than that in the DX structures, consistent with previous reports.^27^ The CD spectra of the structures assembled with polyamines were broadly similar to the controls assembled in Mg^2+^ across all polyamines at these concentrations. An earlier study on DNA tetrahedra reported distortion of B-DNA induced by spermine and spermidine due to shifts in the positive band to lower wavelengths.^16^ However, we did not observe any change relative to the magnesium control. Therefore, our results suggest that at these concentrations, polyamines do not induce any major changes to the B-form of the double helical domains within DX and PX nanostructures.

We also tested the effect of polyamines on the thermal stability of the DNA nanostructures by monitoring the absorbance at 260 nm over a temperature range of 15–90 °C. The melting curves of polyamine-stabilized DX-A resembled those of Mg^2+^ control with a single transition centered around 55–66 °C. DX-P appeared to deviate from a two-state transition model in both control as well as the polyamine-stabilized structures and melts in two stages (**Figure 2e** and **Table S2**). This phenomenon was most prominent in putrescine-assembled DX-P and least prominent in spermine-assembled DX-P. Similar melting behavior is seen other multistranded systems such as DNA triplexes and complexes of bimodal PNAs.^35,36^ All polyamines displayed destabilization with the lower-temperature melting transition, while all except spermine displayed no change in the high-temperature melting transition. It is not clear what these states correspond to, and more studies will reveal whether there are structurally weak regions in DX-P that melts at lower temperature without affecting the rest of the structure and whether there are distinct binding sites for polyamines and Mg^2+^ on the structure affecting their melting behavior.

Overall, polyamines generally reduced the thermal stability of the nanostructure relative to the Mg^2+^ control. We observed minimal impact on the thermal stability of PX and DX-P, with ΔT_m_ values of ∼3.8 °C for PX and ∼4.1 °C for DX-P compared to the structures assembled in Mg^2+^. Thermal stability of DX-A was compromised more by polyamines. All polyamines, except putrescine, decreased the T_m_ by at least 7 °C. Spermine showed the highest destabilizing effect with a ΔT_m_ of ∼12 °C. While putrescine showed the least impact on the thermal stability of DX-A, it was more pronounced on DX-P and PX.

These results suggest that while polyamines may produce nanostructures with high yields, substituting Mg^2+^ with polyamines may compromise the structural stability of the structures. We tested the thermal stability of polyamine-stabilized DNA nanostructures only at one concentration, and any concentration dependent effect on the stability requires further investigation. Nevertheless, the mean T_m_ of all the structures are above 50 °C and therefore multi-crossover structures assembled using polyamines may be suitable for most applications. This could also be useful for applications that involve dynamic reconfiguration of nanostructures where high thermal stability is a liability.

Next, we tested whether polyamines could assist in the formation of motif-based DNA 2D arrays. We chose the AB-type 2D array based on DX motifs^37^ and tested array formation for all the polyamines at concentrations optimized for DX-A (**Figure S4**). The array is self-assembled through two DX tiles A and B*, with tile B* containing a hairpin to observe the periodicity of the array.^37^ Compared to the array in Mg^2+^, we observed that all the polyamines tested yielded 2D arrays with defined striped features but to different extents (**Figure 3**). Putrescine and BAPD yielded well-ordered arrays in hundreds of nanometers whereas spermine, spermidine, and TAEA only formed smaller arrays.

**Figure 3.**
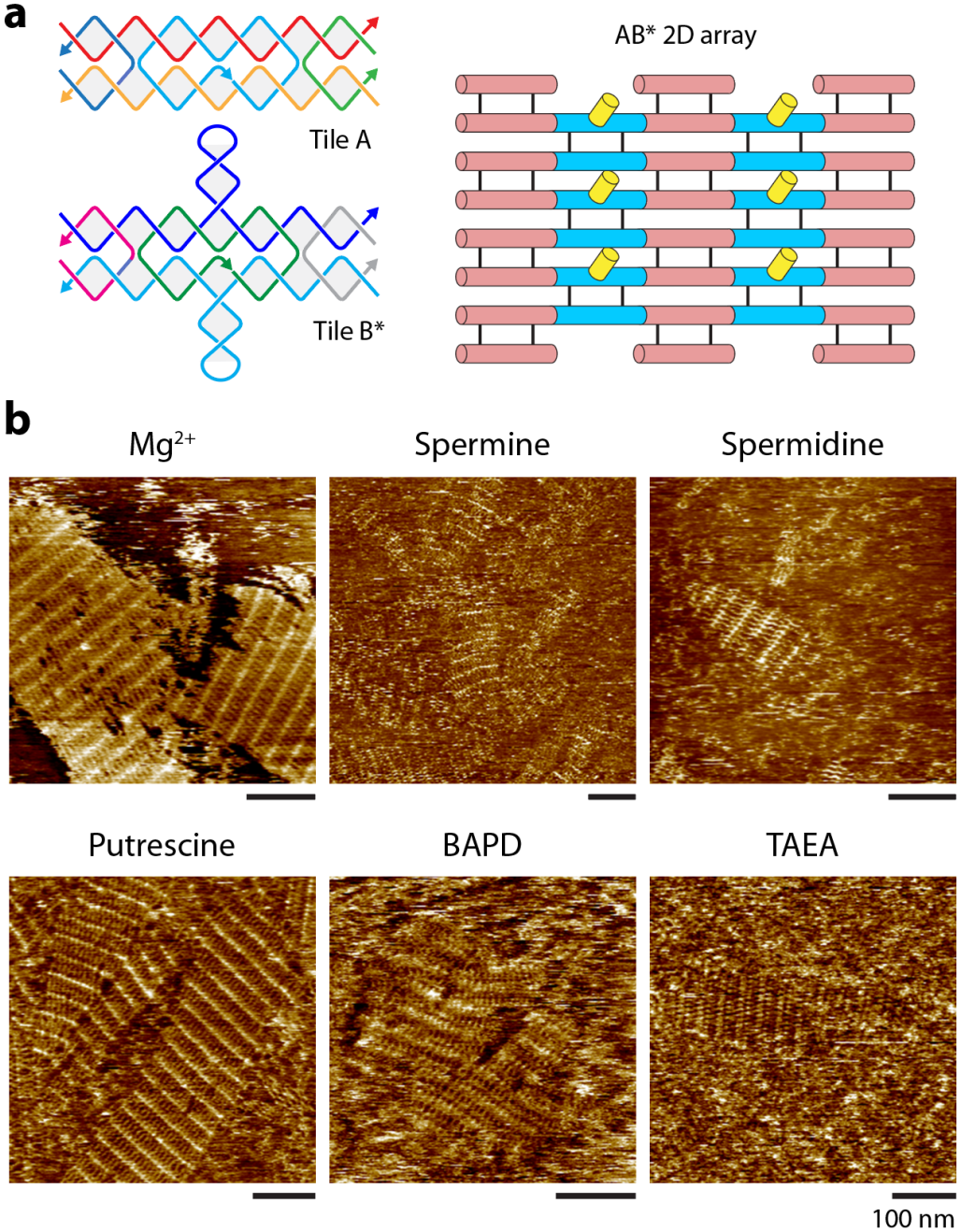
2D self-assembly in polyamines. (a) Schematic of the DX tiles used and the resulting 2D array. (b) AFM images of 2D arrays assembled in magnesium ion and different polyamines.

Our study shows successful assembly of multi-crossover molecules in different polyamines. The concentration of polyamines needed for proper self-assembly depends on the crossover density in the structure, and to some extent, the routing of the strands within the two double helical domains. For these motifs, CD spectra shows that the underlying DNA structure is not affected by polyamines, and while thermal stability is slightly reduced when assembled with polyamines, the structures should still be stable for various applications. While we only compared the polyamines to Mg^2+^ here, prior simulation work showed that spermine binds tightly to helices in DNA origami and reduces the repulsion of parallel helices similar to magnesium while sodium binds weakly and quickly exchanges with the solvent.^38^ In vivo, the concentration of free magnesium ions in the serum is about 0.40–0.68 mM and about 1 mM in the cytoplasm and in the extracellular space,^39,40^ which is inadequate to support DNA nanostructure assembly on their own. Therefore, polyamines can be effective substitutes in this regard to assemble and study muti-crossover DNA nanostructures inside cells since these structures can be assembled under biologically relevant concentrations of polyamines. Our results can also be combined with prior work to create DNA nanoparticle networks^41^ and dynamic DNA nanostructures containing photoresponsive polyamines.^42^ In some cases, specific polyamines can be combined with metal ions such as zinc to achieve reversible condensation of DNA.^43^ A follow up would be to test the effect of polyamines such as spermine on RNA nanostructures since spermine is known to condense DNA but not RNA duplexes.^44^

## Competing interests

The authors have no competing interests.

## Associated content

Materials and methods, additional experimental results and sequences used.

## Author contributions

BRM designed and performed the experiments, analyzed and visualized the data, and wrote the paper. GG, NK, and MZ assembled the nanostructures and characterized them using gel electrophoresis, CD and UV melting experiments. VM performed AFM imaging and analysis. KH provided AFM resources, discussed the results and edited the paper. ARC conceived the project, designed experiments, analyzed and visualized the data, supervised the project and wrote the paper.

## Acknowledgments

Research reported in this publication was supported by the National Institutes of Health (NIH) through National Institute of General Medical Sciences (NIGMS) under award number R35GM150672 (to A.R.C) and R35GM124720 (to K.H).

## Notes

### Competing Interest Statement

The authors have declared no competing interest.

