## Supporting information for "Natural and synthetic polyamines as organic counterions for multi-crossover DNA nanostructures"

**Materials**

DNA strands were purchased from Integrated DNA Technologies with standard desalting. Spermine tetrahydrochloride (99%), Putrescine (99%), N,N'-Bis(3-aminopropyl)ethylenediamine (98%), and Tris(2-aminoethyl)amine (97%) were obtained from Fisher Scientific. Spermidine (>98%) was purchased from Millipore Sigma.

**Preparation of DNA complexes**

DNA double crossover (DX) and paranemic crossover (PX) motifs were prepared by mixing the component oligonucleotide strands stoichiometric ratio in tris-acetate-EDTA buffer containing 40 mM tris base (pH 8), 20 mM acetic acid, and 2 mM EDTA (1× TAE) with either 12.5 mM magnesium acetate or the required concentration of polyamine. The mixture was then annealed using a stepwise annealing protocol on a thermocycler using the following steps: 90°C for 3 min, 65°C for 20 min, 45°C for 20 min, 37°C for 30 min, 20°C for 30 min, then cooled to 4°C. The solutions were incubated at 37 °C for 2 hours instead of thermal annealing for isothermal assembly. The annealed nanostructures were stored at 4°C.

DNA 2D arrays were assembled by annealing 10 strands that constitute the tiles DXT-A and DXT-B* together at an equimolar concentration of 400 nM in 1x TAE buffer supplemented with 12.5 mM magnesium acetate or the required concentration of one of the polyamines. The concentration of spermine, spermidine, putrescine, BAPD, and TAEA were 10 μM, 150 μM, 10 mM, 75 μM, and 75 μM, respectively. Sequences of both the tiles are provided in **Table S1**. Microcentrifuge tubes containing 50 μL of the DNA solution were floated in a 2 L beaker of hot water (~90 °C) and was sealed inside a styrofoam box. The samples were allowed to cool to room temperature gradually over 48 hours.

**Gel electrophoresis**

Non-denaturing gels were prepared using 19:1 acrylamide:bis-acrylamide solution (AccuGel, National Diagnostics). Gels were run using 1× TAE with 12.5 mM $Mg^{2+}$ running buffer at a constant volage at 4 °C. Gels were stained in 0.5× GelRed (Biotium) and imaged using Gel Doc XR+ (BioRad) with standard settings for GelRed illumination and analyzed using ImageLab software. The intensity of the bands was quantified by densitometric analysis. For quantification, the gels were irradiated with UV light for different exposure times and the highest exposure time that did not produce saturation in the bands of interest were used for analysis. The background corrected band intensity of the DX motifs at each polyamine concentration was measured as adjusted volume using Image Lab software. The percentage change in intensity of the band corresponding to samples assembled using $Mg^{2+}$ was treated as relative assembly yield.

### Circular dichroism

DNA motifs were prepared at 2 µM concentration in 1× TAE with either 12.5 mM $Mg^{2+}$ or polyamine at one of the concentrations that produced high relative assembly yield. For DX-A, the concentration of spermine, spermidine, putrescine, BAPD, and TAEA were 10 µM, 150 µM, 10 mM, 75 µM, and 75 µM, respectively. For DX-P, the concentrations of the polyamines were 40 µM, 625 µM, 10 mM, 625 µM, and 625 µM, respectively. For PX, the concentrations were 75 µM, 625 µM, 10 mM, 2.5 mM, and 1.25 mM, respectively. The samples were annealed as described above. 200 µl of the annealed samples was taken in a 1 mm quartz cuvette (Hellma Analytics), and the circular dichroism spectra were recorded using a Jasco-815 CD spectrometer at room temperature. Measurements were taken every 1 nm from 200 nm to 360 nm at a scanning rate of 100 nm/min. Data shown are an average of 3 accumulations.

### UV melting

DX-A, DX-P, and PX motifs were prepared at 1 µM concentration in 1× TAE with 12.5 mM $Mg^{2+}$ or one of the polyamines at specific concentration. The concentration of the polyamine was same as in the CD experiment. Absorbance at 260 nm was recorded from 15°C to 95 °C every 0.5 °C using a Cary 3500 UV-Visible Spectrophotometer (Agilent). The melting curves were normalized 0–1 and fitted to the Boltzmann function using OriginLab software. Melting temperatures were obtained using the first derivative of the fitted curve.

### AFM imaging

The 2D arrays of DXA-DXB* tiles were imaged using Cypher-S Atomic Force Microscope (Asylum Research, Oxford Instruments) with liquid droplet probe holder and BL-AC40TS (Olympus) cantilever in tapping mode (AC mode). 5 µl of annealed DXA-DXB* tiles solution was deposited on a freshly cleaved mica surface and left to stand for 1 min and 100 µl of 1× TAE-$Mg^{2+}$ was added before imaging.

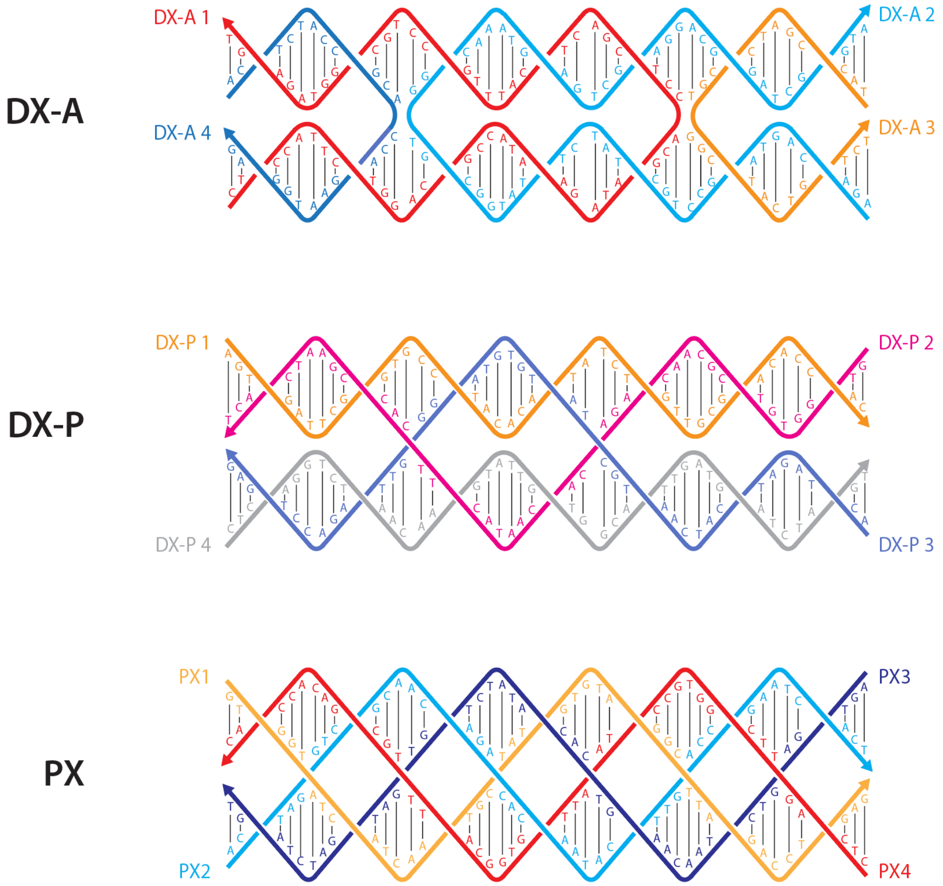

**Figure S1.** Design of motifs with sequences.

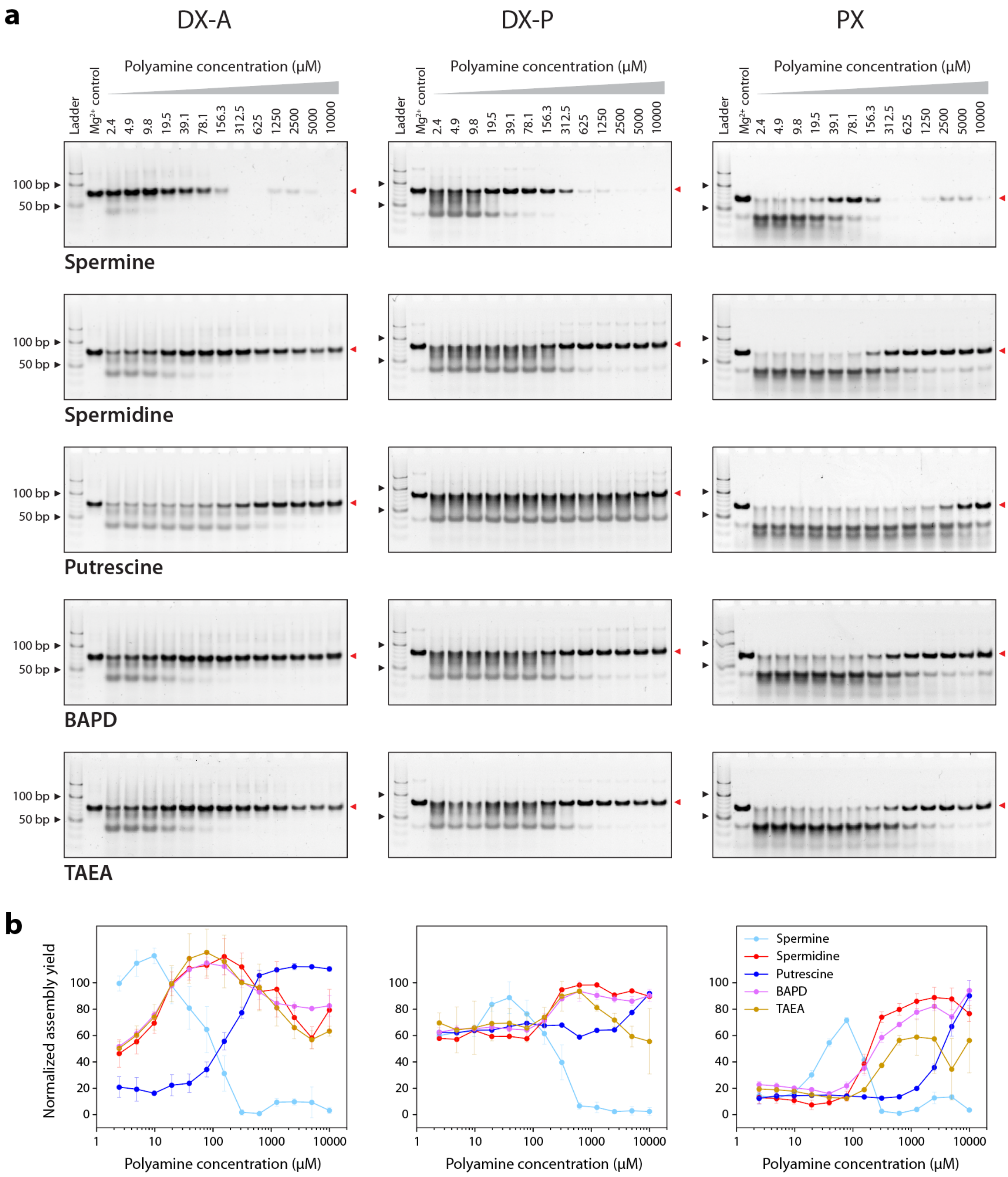

**Figure S2.** (a) Non-denaturing polyacrylamide gels showing the effect of polyamine concentration on the assembly of DX-A, DX-P, and PX. (b) Assembly yields of nanostructures in polyamines. Yields are normalized to the assembly yield in $Mg^{2+}$ containing buffer.

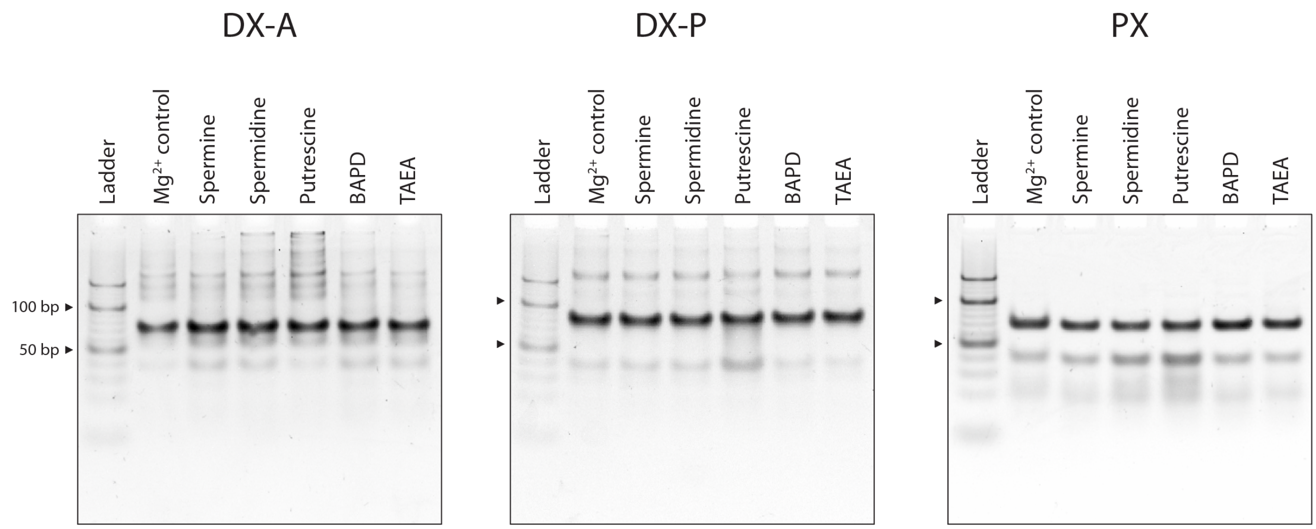

**Figure S3.** Isothermal assembly of motifs at 37 °C.

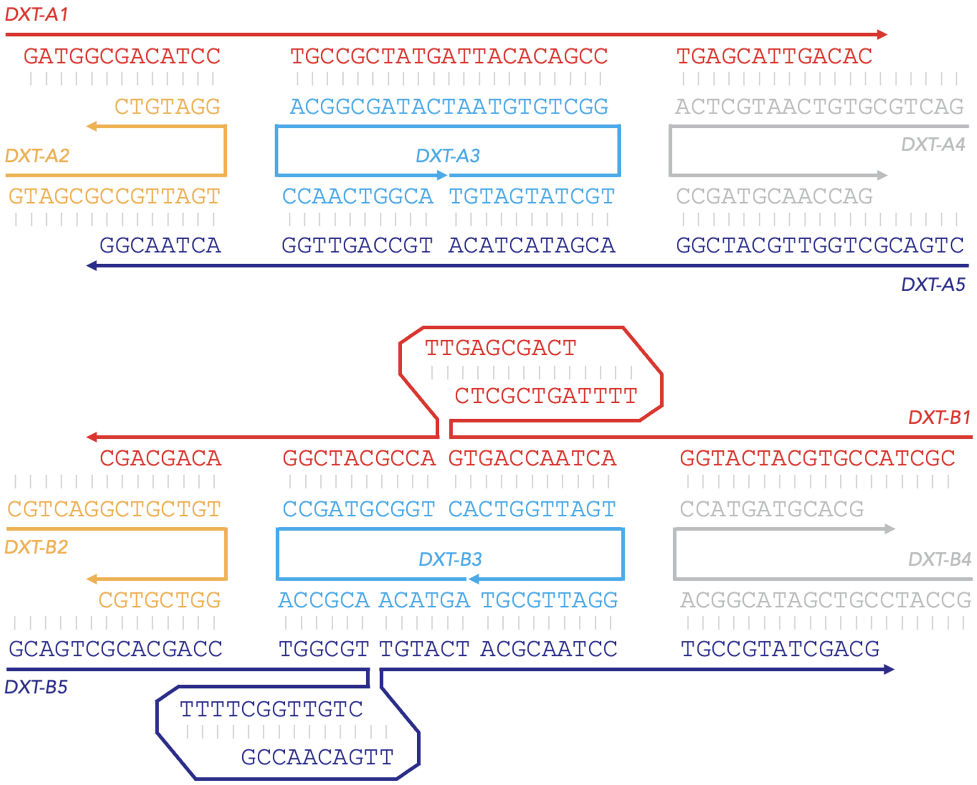

**Figure S4.** Design of motifs used in 2D array.

**Table S1.** Sequences used in this study (written 5′-3′).

| | |
|---|---|
| **Double crossover with antiparallel crossovers (DX-A)** | |
| DXA-1 | CTC CAT TCT GGA CGC CAT AAG ATA GCA CCT CGA CTC ATT TC CG CGG TAG AGT |
| DXA-2 | AGA CAG TAG CCT GCT ATC TTA TGG CGT GGC AAA TGA GTC GAG GAC GGA TCG GTA |
| DXA-3 | ACT CTA CCG CAC CAG AAT GGA G |
| DXA-4 | TAC CGA TCC GTG GCT ACT GTC T |
| **Double crossover motif with parallel crossovers (DX-P)** | |
| DXP-1 | AGT GAT TCG GTG CCT ACA CAT AT TGT TC GAC ACC AC |
| DXP-2 | GTG GG TCG CAA CAG ACA CAA TAC TTC ACC GAA TCA CT |
| DXP-3 | ACT AGA TCA TCA ATG CTA TGT GTA GGG TTA GAC CTG AG |
| DXP-4 | CTC AGG TCT AAC AAG TAT TGT GGC ATT GAT GAT CTA GT |
| **Paranemic crossover (PX) motif** | |
| PX-1 | GTG GTA TCA TCA ATG CTA TGT GTA GGC TTA GAC CG AG |
| PX-2 | ACT AGG TCG CAA CAG ACA CAA TAC TTG ACC GAA TA CT |
| PX-3 | AGT GAG TCT AAC AAG TCA CAT ATC TGT GAT GAT CA GT |
| PX-4 | CTC AGT TCG GTG CCT AAT TGT GGC ATT TGC GAC ACC AC |
| **DX tile A for 2D array** | |
| DXT-A1 | GAT GGCGAC ATC CTG CCG CTA TGA TTA CAC AGC CCT GAG CAT TGA CAC |
| DXT-A2 | GTA GCG CCG TTA GTG GAT GTC |
| DXT-A3 | TGT AGT ATC GTG GCT GTG TAA TCA TAG CGG CA |
| DXT-A4 | GAC TGC GTG TCA ATG CTC ACC GAT GCA ACC AG |
| DXT-A5 | CTG ACG CTG GTT GCA TCG GAC GAT ACT ACA TGC CAG TTG GAC TAA CGG |
| **DX tile B for 2D array** | |
| DXT-B1 | CGC TAC CGT GCA TCA TGG ACT AAC CAG TGC TCG CTG ATT TTT CAG CGA GTT ACC GCA TCG GAC AGC AGC |
| DXT-B2 | CGT CAG GCT GCT GTG GTC GTG C |
| DXT-B3 | AGT ACA ACG CCA CCG ATG CGG TCA CTG GTT AGT GGA TTG CGT |
| DXT-B4 | GCC ATC CGT CGA TAC GGC ACC ATG ATG CAC G |
| DXT-B5 | GCA GTC GCA CGA CCT GGC GTC TGT TGG CTT TTG CCA ACA GTT TGT ACT ACG CAA TCC TGC CGT ATC GAC G |

**Table S2.** Melting temperature ($T_m$) of DNA nanostructures assembled using polyamines

| Nanostructure | Counterion | $T_m$ (°C) | $\Delta T_m$ (°C) |
|---|---|---|---|
| DX-A | $Mg^{2+}$ | 67.3 | |
| | Spermine | 55.0 | -12.3 |
| | Spermidine | 59.7 | -7.6 |
| | Putrescine | 66.4 | -0.9 |
| | BAPD | 56.4 | -10.9 |
| | TAEA | 56.2 | -11.1 |
| DX-P | $Mg^{2+}$ | 63.4 | |
| | Spermine | 58.3 | -5.1 |
| | Spermidine | 62.0 | -1.4 |
| | Putrescine | 55.5 | -7.9 |
| | BAPD | 60.8 | -2.6 |
| | TAEA | 60.0 | -3.4 |
| PX | $Mg^{2+}$ | 58.2 | |
| | Spermine | 55.1 | -3.1 |
| | Spermidine | 53.8 | -4.4 |
| | Putrescine | 52.0 | -6.2 |
| | BAPD | 55.3 | -2.9 |
| | TAEA | 55.6 | -2.6 |
